# ASF1 activation PI3K/AKT pathway regulates sexual and asexual development in filamentous ascomycete

**DOI:** 10.1101/2021.10.18.464864

**Authors:** Shi Wang, Xiaoman Liu, Chenlin Xiong, Susu Gao, Wenmeng Xu, Lili Zhao, Chunyan Song, Zhuang Li, Xiuguo Zhang

**Author notes:** Correspondence and requests for materials should be addressed to Z.L & X.G.Z,).

## Abstract

Sexual and asexual reproduction is ubiquitous in eukaryotes. PI3K/AKT signaling pathway can modulate sexual reproduction in mammals. However, this signaling pathway modulating sexual and asexual reproduction in fungi is scarcely understood. SeASF1, a SeH4 chaperone, could manipulate sexual and asexual reproduction of *Stemphylium eturmiunum*. SeDJ-1, screened from SeΔ*asf1* transcriptome, was confirmed to regulate sexual and asexual development by RNAi, of which the mechanism was demonstrated by detecting transcriptional levels and protein interactions of SeASF1, SeH4 and SeDJ-1 by qRT-PCR, and Y2H, Co-IP and Pull-down, respectively. SeASF1 coupling SeH4 bound SeDJ-1 to arouse the sexual and asexual activity. In *S. eturmiunum* genome, SeDJ-1 was upstream while SeGSK3 was downstream in PI3K/AKT signaling pathway. Moreover, SeDJ-1 interacted with SePI3K or SeGSK3 in *vivo* and in *vitro*. Significantly, SeDJ-1 or SePI3K could effectively stimulate sexual activity alone, but SePI3K could recover the sexual development of SiSeDJ-1. Meanwhile, SeDJ-1-M6 was a critical segment for interaction of SeDJ-1 with SePI3K. SeDJ-1-M6 played a critical role in irritating sexual reproduction in SiSePI3K, which further uncovered the regulated mechanism of SeDJ-1. Summarily, SeASF1 coupling SeH4 motivates SeDJ-1 to arouse SePI3K involved in sexual reproduction. Thus, SeASF1 can activate PI3K/AKT signaling pathway to regulate sexual and asexual development in filamentous ascomycete.

## Introduction

Sexual reproduction is the predominant reproductive strategy in eukaryotes (Dacks and Roger, 1999; Ramesh et al., 2005). A series of factors including mating-types (Böhm et al., 2013; Coppin et al., 1997), pheromone components (Bobrowicz et al., 2002; Lin et al., 2011), G proteins (Li et al., 2007; Studt et al., 2013) and velvet proteins (Bayram and Braus, 2012) are involved in sexual reproduction in fungi. MAPK (Mitogen-Activated Protein Kinase) (Bayram et al., 2012; Chen et al., 2002; Saito, 2010), CWI (Cell Wall Integrity) (Teichert et al., 2014; Zhang et al., 2020) or cAMP-PKA (cyclic Adenosine Monophosphate/Protein Kinase A) signaling pathway1 (Dos Reis et al., 2019) is a highly conserved signaling cascade in eukaryotes and is also required for sexual mating in fungi. More than 20 hypotheses have been used to reveal why sexual reproduction is maintained in fungi (de Visser and Elena, 2007; Hadany and Comeron, 2008). However, most of them devoted to maintenance of fungi sexual mating were still remained unclearly. Thus, sexuality in fungi becomes more perplexing yet intriguing facets of biology that is inevitable to breed a series of magical mechanisms.

ASF1 is first identified in *Saccharomyces cerevisiae* and carries out an important role for mating type (Le et al., 1997). ASF1, a H3-H4 chaperone, is highly conserved from yeast to mammals and involved in nucleosome assembly/disassembly (Avvakumov et al., 2011; Eitoku et al., 2008; Prado et al., 2004; Min et al., 2020; Sanematsu et al., 2006), normal cell cycle progression (Groth et al., 2007, Sutton et al., 2001), genomic instability along with histone modification (Das et al., 2014; Li et al., 2008; Recht et al., 2006; Yuan et al., 2009), DNA replication, repair, recombination and transcriptional regulation (Mousson et al., 2007). ASF1 has a serious of magic for modulating female reproduction in mice (Messiaen et al., 2016) and requiring for heat stress response and gametogenesis in *Arabidopsis thaliana* (Zhu et al., 2011; Weng et al., 2014; Min et al., 2019). Meanwhile, ASF1 can also manipulate the sexual reproduction in *Sordaria macrospora* effectively (Gesing et al., 2012). Summarily, ASF1 shows a ubiquitous function for regulating sexual development in animals, plants and fungi. However, whether ASF1 can activate a signaling pathway to mediate sexual or asexual development in filamentous ascomycete is barely accepted.

DJ-1, named as PARK7, is first reported to associated with Parkinson’s disease (PD) (Bonifati et al. 2012) and then verifies to be an oncogene for mediating the regulation of numerous types of cancer (Bai et al., 2012; Chen et al., 2012; Scumaci et al., 2020). DJ-1 is an essential regulator of multiple cellular processes, including anti-oxidative stress, anti-apoptotic effects, and protein degradation (Taira et al., 2004; Mukherjee et al., 2015; Hijioka et al., 2017). As a multifunctional protein, DJ-1 plays a major role in the phosphatidylinositol-3-kinase (PI3K)/AKT signaling pathway (Yang et al., 2005). The PI3K/AKT signaling pathway manipulates a variety of biological processes, including cell differentiation, proliferation, growth, metabolism, survival, genomic stability, protein synthesis, angiogenesis, cancer (Yang et al., 2018; Engelman et al., 2006; Liu et al., 2020; Patra et al., 2019; Zhou et al., 2017), and even inhibition of apoptosis and oxidative stress and regulation of a variety of downstream molecules (Srinivasan et al., 2005; Gong et al., 2018). Significantly, PI3K/AKT signaling pathway can modulate sexual reproduction in mammals (Shao et al., 2019; Fu et al., 2020). However, the role of DJ-1 or ASF1 in connection with DJ-1 mediates this signaling pathway to irritate sexual and asexual reproduction in filamentous ascomycete is hardly understood.

*Stemphylium* and its two closely related genera *Alternaria* and *Ulocladium* belong to filamentous ascomycetes (Simmons., 1967). *Stemphylium* was subject to the ascomycete family Pleosporales and Pleosporaceae (Câmara et al., 2002; Simmons, 1989). The sexual states of *Stemphylium* and *Alternaria* are *Pleospora* and *Lewia*, respectively (Lucas and Webster, 1964; Simmons, 1969, 1989; Inderbitzin et al., 2005), but the sexual state of *Ulocladium* has no yet been identified (Wang et al., 2017). Most species within these three similar genera are mainly allied to asexual states (Woudenberg et al., 2013, 2017; Câmara et al., 2002). The taxonomic study of them are mainly focused on asexual means. Until now, very few asexual species of them are corresponding to sexual type, which is still challenged due to lack of understanding the mechanisms of these states.

In this study, we investigated the biological function of SeASF1 for regulating sexual and asexual development in *S. eturmiunum*. We showed that SeASF1, identified from *S. eturmiunum*, could modulate sexual development in *S. macrospora* by heterologous expression analysis and was equipped to activate sexual and asexual reproduction in *S. eturmiunum*. A variety of up-regulated or down-regulated genes, including Se02026 (SeDJ-1, unidentified protein), Se01950 (Heat shock protein), Se03485 (LysM domain-containing protein), Se04320 (Proline dehydrogenase), Se07693 (vesicle coat complex COPII, subunit SEC31), Se10206 (Allantoate permease) and Se10302 (Choline dehydrogenase), were identified by comparative analysis of transcriptome data. As a result, Se02026 (SeDJ-1) exhibited a unique characteristic for carrying out sexual and asexual development of *S. eturmiunum* by gene silencing. Subsequently, our experiments demonstrated that SeDJ-1 could directly interact with SeASF1 and SeH4 or with SePI3K and SeGSK3 in PI3K/AKT pathway in *vivo* and in *vitro*, respectively. Furthermore, seven truncations of *Sedj-1* (*Sedj-1*-M1, *Sedj-1*-M2, *Sedj-1*-M3, *Sedj-1*-M4, *Sedj-1*-M5, *Sedj-1*-M6 and *Sedj-1*-M7) were obtained and confirmed that *Sedj-1*-M6 was a key domain for modulating *Sedj-1* interaction with SePI3K. At the same time, *Sedj-1*-M6 was further confirmed to play a crucial role in triggering SePI3K to modulate sexual features contrast to *Sedj-1*-M7. Meanwhile, our study verified that *Sepi3k* could motivate sexual reproduction in Si*Sedj-1* strains reversely. In summary, SeASF1 could bind with SeDJ-1 to irritate SePI3K for modulating sexual and asexual reproduction. Thus, PI3K/AKT pathway is assumed to involve in sexual and asexual development in filamentous ascomycetes.

## Results

### Identification and characterization of ASF1 gene in *S. eturmiunum*

ASF1 was identified from *S. eturmiunum* genome database (unpublished) by PCR amplification. Primers for PCR were designed by Primer express 3.0 software (Supplemental Table S2). This gene was named as *Seasf1* (KX033515). SeASF1 has 291 amino acids with a calculated molecular mass of 31.98 kDa. Alignment of SeASF1 sequence with its homologous sequences from plants, animals and other fungi species (https://www.ncbi.nlm.nih.gov/) (Supplemental Table S1) revealed that the N-terminus sequences (1-154 aa) of ASF1 was highly conserved and contained the ASF1 hist chap superfamily functional domain, but the C-terminus was diversity (Supplemental Figure S2). Phylogenetically, all these analyzed sequences were grouped into three clusters that were labeled by different background colors. The cluster1 was divided into three subclusters with different background colors. Also, SeASF1 shared 98.28% similarity with ASF1 of *S. lycopersici* (KNG47682), and 90.51% to 91.13% similarity with ASF1 of two *Pyrenophora* species (XP_003295297 and XP_001936595). Notably, SeASF1 shared 92.78% identify with ASF1 of *Setosphaeria turcica* (XP_008022755), and more than 91% identify with ASF1 of five *Bipolaris* species (XP_007710420, XP_007695597, XP_014075349, XP_014554985 and XP_007689798), respectively. However, SeASF1 shared 45.45% similarity with ASF1 of *Schizosaccharomyces pombe* (CAA20365) (Supplemental Figure S1). All these data indicate that ASF1 is widely distributed in eukaryotic organisms.

### SeASF1 restores the phenotype of sexual reproduction in SmΔ*asf1* mutant

In previous study, the *S. macrospora* ASF1 (XP003345657) was localized to the nucleus and essential for sexual reproduction (Gesing et al., 2012). Alignment of SeASF1 sequence with its homologous sequences from plants, animals, and other fungi species showed that ASF1 has a specifically conserved function domain (Supplemental Figure S2), we doubted whether *Seasf1* operates a similar conservatively function. Here, we heterologously expressed the *S. eturmiunum asf1* in the SmΔ*asf1* mutant, and obtained two heterologous expression transformants which were succeeded in complementing the developmental defects of SmΔ*asf1* strain (Supplemental Figure S3A). Two transformants, SmΔ*asf1*::GFP*Seasf1*-1/2, were expressed by fusion constructs that were identified by PCR and western blot, respectively (Supplemental Figure S4) (The primal PCR result is shown in Supplemental Figure S23, and the primal western blots results are shown in Supplemental Figure S24, S25). Sexual reproduction of these two transformants was completed after growing on CM medium for 10 days, and perithecia carrying mature asci after inducing for 20 days (Supplemental Figure S3B). Fluorescence microscope showed that *Seasf1* was also localized to the nucleus in *S. macrospore* (Supplemental Figure S3C). Taken together, heterologous expression analyses verify that SeASF1 has a conserved function, as well as SmASF1, for producing sexual reproduction in filamentous fungi.

### SeASF1 modulates asexual and sexual development in *S. eturmiunum*

To uncover the biological functions of the SeASF1 during vegetative and sexual development of *S. eturmiunum*, we obtained two SeΔ*asf1* mutant strains: SeΔ*asf1*-0::EGFP and SeΔ*asf1*-5::EGFP, and two complemented transformants: SeΔ*asf1*-0::EGFP*Seasf1* and SeΔ*asf1*-5::EGFP*Seasf1*. Two knockout mutants were detected by southern blot and PCR (Supplemental Figure S5B, C) (The primal PCR results are shown in Supplemental Figure S26, S27, and the primal southern blot result is shown in Supplemental Figure S28). Two complemented transformants were also detected using western blot and PCR (Supplemental Figure S6) (The primal PCR result is shown in Supplemental Figure S29, and the primal western blots results are shown in Supplemental Figure S30, S31). To determine the role of *Seasf1* in hyphal and colonial growth, these four mutants and WT strains were inoculated on PDA medium for 9 days, respectively. The cultures were then photographed after 1 day, 3 days, 5 days, 7 days and 9 days (Supplemental Figure S7A). In comparison to WT, two complemented transformants were almost returned to normal growth as well as colonial and hyphal growth (Supplemental Figure S7B, C). However, two SeΔ*asf1* strains produced the hyphal fusion, anomalously distributed of nucleus in mycelium, and abnormal conidia which were significantly opposed to those of the complemented transformants and WT strains (Figure 1A-E). These results suggest that *Seasf1* is involved in asexual development of *S. eturmiunum*. On the other hand, microcosmic observations sexual developmental of these four mutants contrast to WT strains showed two complemented transformants and WT strains produced abundant perithecia and normal asci cultured on CM medium after 4 weeks (Figure 1F). Conversely, SeΔ*asf1* strains were completely sterile and did not produce perithecia and asci under the same condition (Figure 1G). Furthermore, the expression levels of genes, including mainly Gα subunit, MAT1, MAT2, Ste2 and Ste3 involved in MAPK pathway for modulating sexual reproduction, did not change significantly in the two SeΔ*asf1* mutants compared to WT strain (Supplemental Figure S8). These results suggested that SeASF1 might mobilize a new pathway to regulate sexual reproduction in *S. eturmiunum*.

**Figure 1.**
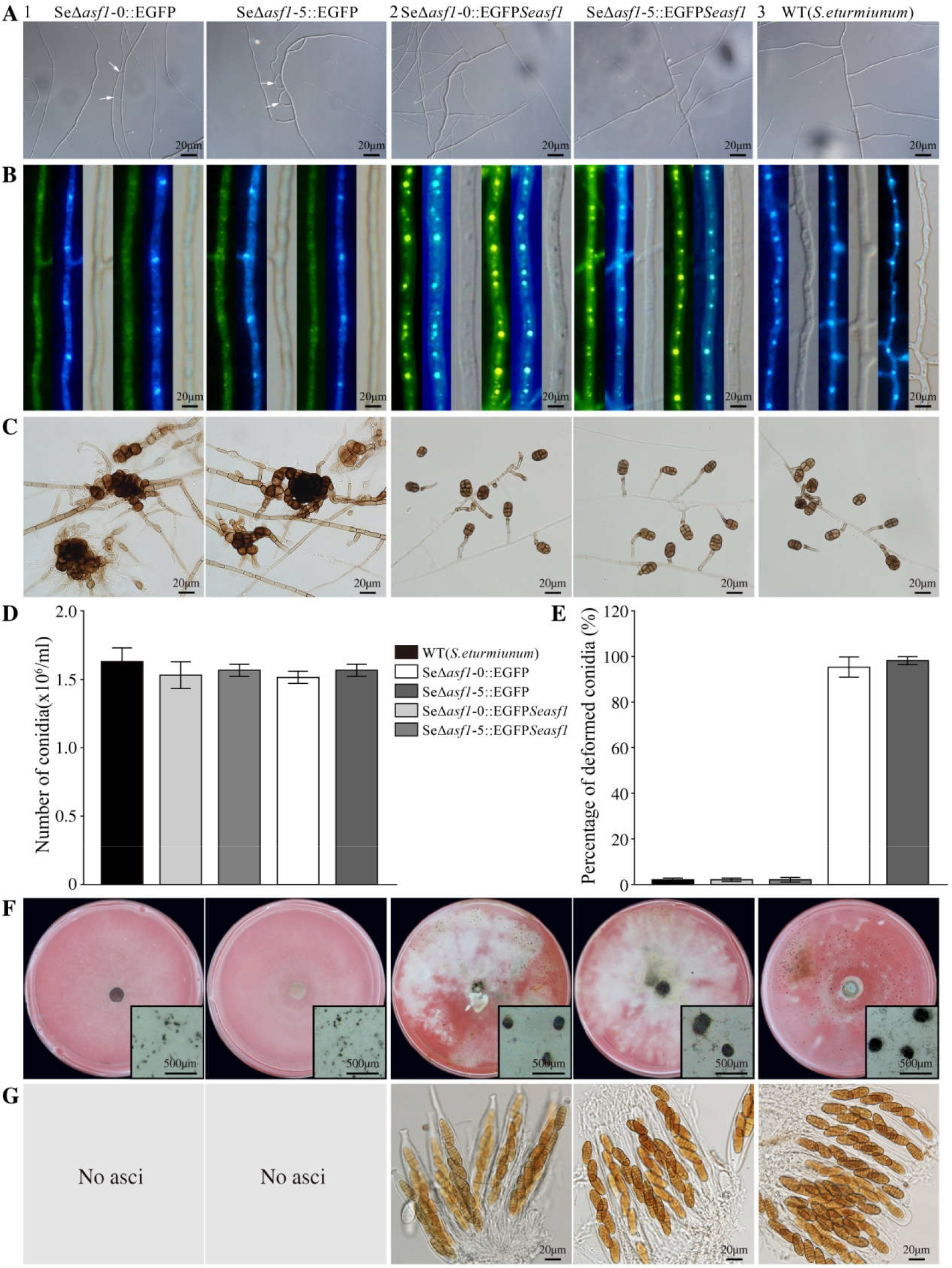
SeASF1 regulates asexual and sexual developmental characterization in *S. eturmiunum*. **A**, Characterizations of hyphal fusion in two SeΔ*asf1* mutants, two SeΔ*asf1*::EGFP*Seasf1* transformants, and WT strains. The images were photographed after growing on PDA medium for 7 days. The fusions in the hyphae were marked with white arrows. **B**, The mycelia of four mutants and WT strains were examined by DIC and fluorescence microscopy for GFP and DAPI after growing on PDA medium for 12 days. **C**, Conidia morphology of four mutants and WT strains were cultured on CM medium for 4 weeks. Bar= 20 μm. **D**, The number of conidia was counted by blood counting chamber. **E**, Percentage of deformed conidia from two SeΔasf1 mutants incubated in CM medium at 4 weeks compared with two SeΔ*asf1*::EGFP*Seasf1* mutants and WT strains. **F** and **G**, Perithecia of four mutants and WT strains were visualized as black structures on CM medium. Perithecia were shown in the lower right. **G**, Two SeΔ*asf1* mutants were not able to produce perithecia that were opposed to two SeΔ*asf1*::EGFP*Seasf1* mutants and WT strains. Photographs were taken at 6 weeks after sexual induction. Bar= 500 μm.

### RNA-seq analysis the differentially expressed genes (DEGs) involved in regulation of the developmental functions

The previous results show that *Seasf1* can modulate asexual and sexual development in *S. eturmiunum* and *S. macrospora*. To further search whether other genes are likely to involve in these developmental functions modulating by *Seasf1*, a comparative analysis of genes expression differences was carried out SeΔ*asf1* and WT-sexual, SeΔ*asf1* and WT-vegetative, and WT-sexual and WT-vegetative. As a result, 3716 DEGs between SeΔ*asf1* and WT-sexual, of which 2342 genes up-regulated and 1374 genes down-regulated. 3023 DEGs between SeΔ*asf1* and WT-vegetative, of which 1719 up-regulated and 1304 down-regulated. 3094 DEGs between WT-sexual and WT-vegetative, of which 1343 up-regulated and 1751 down-regulated (Supplemental Figure S9B). A total of 380 DEGs among three transcripts were identified (fold change >2.0, *q*-value <0.005) (Supplemental Figure S9A) and subsequently analyzed by hierarchical clustering (Supplemental Figure S10). Through the comparative analysis of transcriptome data, we speculated that these significantly up-regulated or down-regulated genes might involve in the SeASF1 regulation of sexual and asexual development.

To further determine the roles of these up-regulation or down-regulation genes, the histograms of GO enrichment analysis of DEGs are depended on the data of SeΔ*asf1* and WT-vegetative, and SeΔ*asf1* and WT-sexual. GO analyses found that a large number of genes are potentially involved in the process of cellular, secondary metabolites and development, cell part and catalytic activity in the development of *S. eturmiunum* (Supplemental Figure S9C, D). Meanwhile, another seven genes including Se02026 (SeDJ-1, unidentified protein), Se01950 (Heat shock protein), Se03485 (LysM domain-containing protein), Se04320 (Proline dehydrogenase), Se07693 (vesicle coat complex COPII, subunit SEC31), Se10206 (Allantoate permease) and Se10302 (Choline dehydrogenase) were predicted to be involved in *Seasf1* practice on asexual and sexual development (Supplemental Table S4). These findings suggest that SeASF1 is likely to confront with other genes to regulate the asexual and sexual development.

### SeDJ-1 plays a role in asexual and sexual development

DJ-1 (SeDJ-1) was cloned from *S. eturmiunum*. SeDJ-1 contains 257 amino acids with a calculated molecular mass of 28 kDa. Phylogenetically, SeDJ-1 sequence with its homologous sequences from plants, animals and other fungi species (https://www.ncbi.nlm.nih.gov/) (Supplemental Table S5) were grouped into three clusters that were labeled by different background colors. The cluster3 contained all of the DJ-1 sequences from multiple fungi species that was divided into three subclusters. Also, SeDJ-1 shared 93.77% similarity with DJ-1 of *S. lycopersici* (RAR14805), and less than 20% similarity with DJ-1 of all other fungi species (Supplemental Figure S11). Thus, DJ-1 is widely distributed in eukaryote, but SeDJ-1 is considerably conserved with DJ-1 of *S. lycopersici*.

To further investigate whether each of these selected seven genes are involved in asexual and sexual development in *S. eturmiunum*, we generated two silenced transformants of each gene by *A. tumefaciens* mediated method. Our experiments confirmed that these seven genes excluded Se02026 (SeDJ-1) compromised on asexual and sexual development (Supplemental Figure S13-S18). Two *Sedj-1*-silenced transformants (Si*Sedj-1*-T1 and Si*Sedj-1*-T4) appeared the slow growth rate of colonies related to the control strains (Supplemental Figure S12A, B). Notably, the nuclei were anomalously distributed in mycelia of two silenced transformants (Figure 2A). Conidiogenous cells of the two silenced lines were swollen at the apex or lateral branch, while they had grown to be secondary mycelia at apex or side in the control strains at 7 days. At 13 days, conidiophores and conidiogenous cells were obvious development and paled in two silenced lines, but conidiogenous cells of the control strains appeared swollen at the apex and darkened and conidia were imaged young, solitary, body brown and ellipsoid to cylindrical. At 20 days, subglobose and young conidia were pictured in two silenced lines, however, the control strains had produced near mature oblong conidia. For two silenced lines, the mature conidia were not discovered but conidiophores were turned into bead-like which were significantly different from control strains at 30 days (Figure 2B). By contrast, the young and irregular ascogonia were produced in the two silenced strains, while the young protoperithecia could be discovered in the control strains at 13 days. At 25 days, however, immature perithecia did not observe in the two silenced strains which were significantly different from the control strains. Moreover, at 34 days, the two silenced strains did not produce the perithecia, but the near mature perithecia with asci had produced in the control strains. Finally, the mature asci were only pictured in control strains at 45 days (Figure 2C). These results indicate that SeDJ-1 plays a crucial role in asexual and sexual development for *S. eturmiunum*.

**Figure 2.**
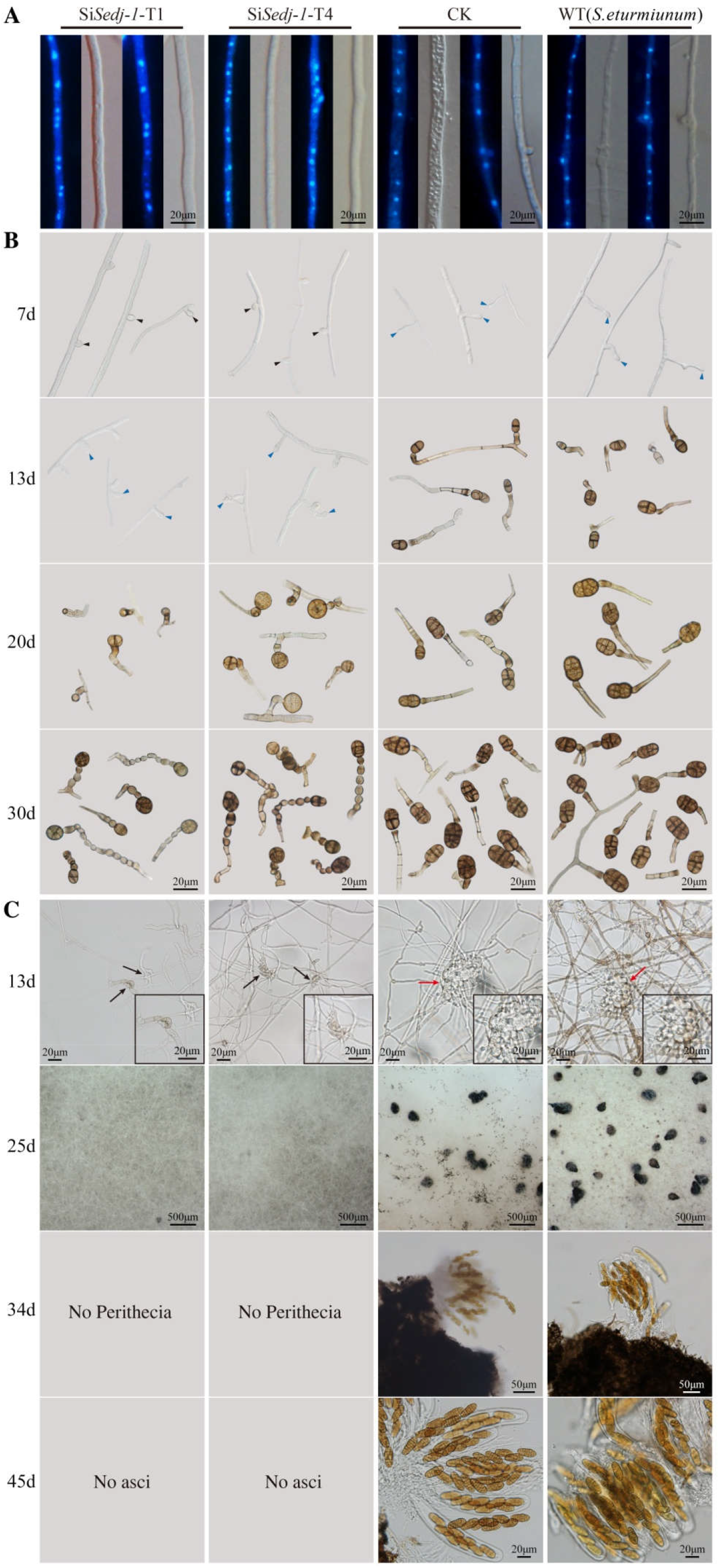
SeDJ-1 plays a role in asexual and sexual development of *S. eturmiunum*. **A**, The mycelium of two silenced transformants, CK and WT strains were incubated on PDA medium for 6 days and examined by DIC and fluorescence microscopy. Two silenced transformants were Si*Sedj-1*-T1 and Si*Sedj-1*-T4. *S. eturmiunum* and control strains were used as WT and CK, respectively. The nuclei of the mycelia were discovered under the fluorescence microscopy after staining by DAPI. **B**, For the microscopic investigation of conidiophores, conidiogenous cells and conidia development, two silenced transformants, CK and WT strains were grown on CM medium for 7 days, 13 days, 20 days and 30 days, respectively. Black arrowheads indicated conidiogenous cells, and blue arrowheads indicated secondary mycelia. **C**, For the microscopic investigation of ascogonia, protoperithecia, young perithecia and asci development, all strains were grown on PDA medium and examined after growth at 25 °C for 13 days, 25 days, 34 days and 45 days, respectively. Insets showed enlarged ascogonia and protoperithecia on the bottom right sides. Perithecia were visualized as black structures. Black arrows indicated ascogonia, and red arrows indicate protoperithecia. Bar= 20 μm, 50 μm and 500 μm.

### SeASF1 interaction with SeH4 or SeDJ-1, and SeH4 interaction with SeDJ-1

To verify whether can occur the interaction between SeASF1 and SeDJ-1, SeDJ-1 and SeH4. Firstly, the transcript levels of *Sedj-1* and *SeH4* were detected in two SeΔ*asf1* mutants. The expressions of *Sedj-1* and *Seasf1* were measured in two Si*SeH4* lines, and those of *Seasf1* and *SeH4* examined in two Si*Sedj-1* lines. As a result, *Sedj-1* and *SeH4* showed down and up-regulation in two SeΔ*asf1* mutants, respectively (Figure 3a). At the same time, *Sedj-1* and *Seasf1* displayed down and up-regulation in two Si*SeH4* lines, respectively (Figure 3B), while *Seasf1* and *SeH4* showed down-regulation in two Si*Sedj-1* lines, respectively (Figure 3C). Therefore, SeDJ-1 was a positive factor for SeASF1 expression. Secondly, Y2H revealed that SeASF1 interacted with SeH4 and SeDJ-1, and SeH4 interacted with SeDJ-1 (Supplemental Figure S19A, B). On the basis of GST pull-down, SeASF1 was specifically interacted with SeH4 (Figure 3D) (The primal western blots of input results are shown in Supplemental Figure S32A, S33A and S34A, while the primal western blots of Pull-down results are shown in Supplemental Figure S32B, S33B and S34B), while SeDJ-1 could interact with SeASF1 and SeH4, respectively (Figure 3E) (The primal western blots of input results are shown in Supplemental Figure S35A, S36A and S37A, while the primal western blots of Pull-down results are shown in Supplemental Figure S35B, S36B and S37B). Finally, all those results of the pull-down experiments were further assured by Co-IP assays (Figure 3F, G) (f-left: The primal western blots of input results are shown in Supplemental Figure S38A, S39A and S40A, while the primal western blots of IP results are shown in Supplemental Figure S38B, S39B. f-right: The primal western blots of input results are shown in Supplemental Figure S41A, S42A and S43A, while the primal western blots of IP results are shown in Supplemental Figure S41B, S42B) (g-left: The primal western blots of input results are shown in Supplemental Figure S44A, S45A and S46A, while the primal western blots of IP results are shown in Supplemental Figure S44B, S45B. g-right: The primal western blots of input results are shown in Supplemental Figure S47A, S48A and S49A, while the primal western blots of IP results are shown in Supplemental Figure S47B, S48B). Thus, SeASF1 interacted with SeH4 or SeDJ-1, and SeH4 also interacted with SeDJ-1. Altogether, SeDJ-1 could cooperate with SeASF1 and SeH4 to modulate asexual and sexual development of *S. eturmiunum*.

**Figure 3.**
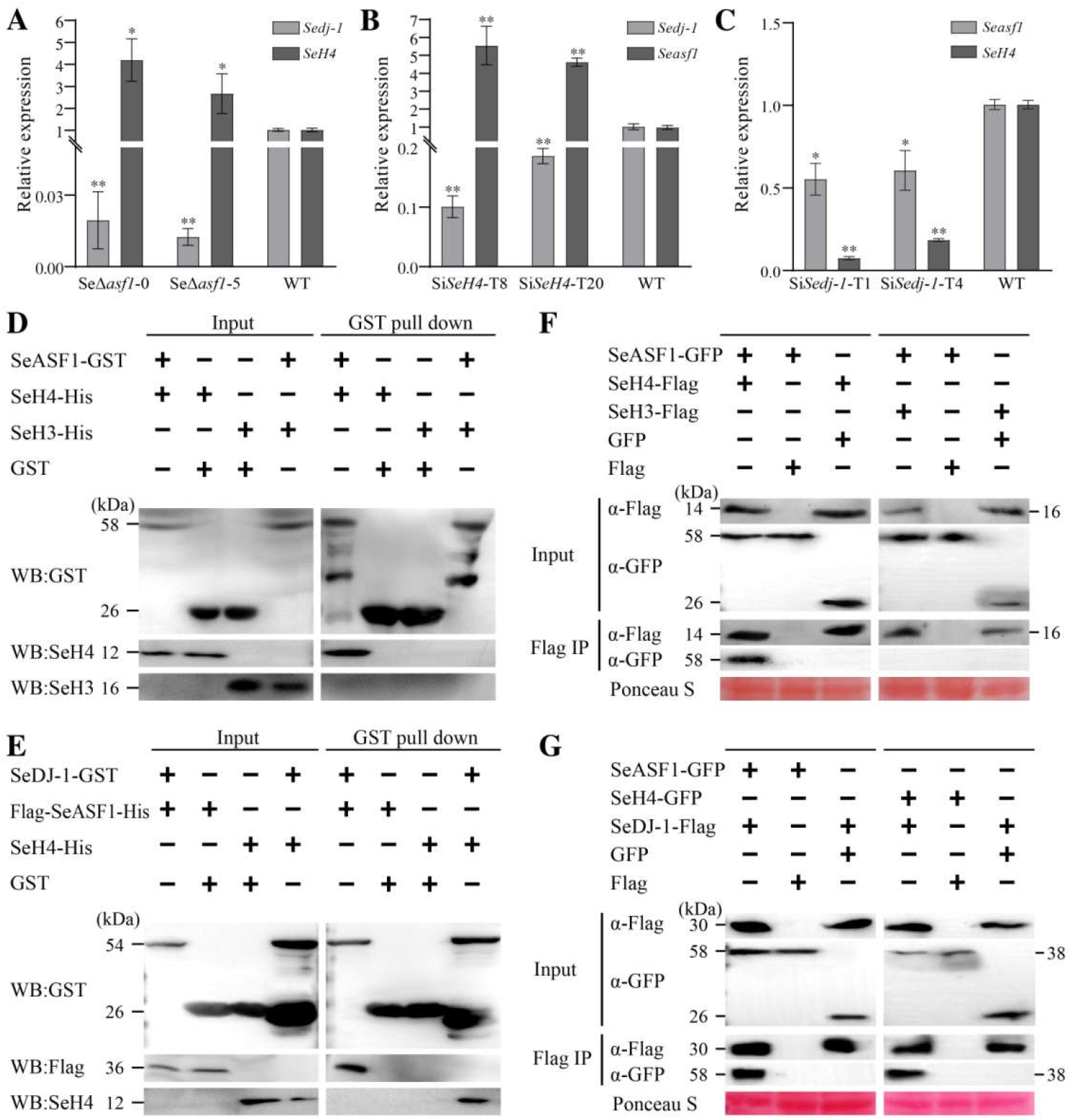
SeASF1 interaction with SeH4 or SeDJ-1, and SeH4 interaction with SeDJ-1. **A**, The expression levels of *Sedj-1* and *SeH4* in two SeΔ*asf1* mutants were measured by qRT-PCR. **B**, The expression levels of *Sedj-1* and *Seasf1* in two Si*SeH4* lines were measured by qRT-PCR. **C**, The expression levels of *Seasf1* and *SeH4* in two Si*Sedj-1* lines were measured by qRT-PCR. The degree of WT was assigned to value 1.0. Two *Seasf1* deleted mutants were SeΔ*asf1*-0 and SeΔ*asf1*-5. Two *SeH4*-silenced lines were Si*SeH4*-T8 and Si*SeH4*-T20. Two *Sedj-1*-silenced lines were Si*Sedj-1*-T1 and Si*Sedj-1*-T4. *S. eturmiunum Actin* was used as endogenous control. The bars indicated statistically significant differences (ANOVA; \**P* < 0.05, \*\**P* < 0.01). **D**, SeASF1 was cloned into plasmid pGEX-6P-1. SeH4 or SeH3 was cloned into plasmid pET28a. SeASF1-GST was expressed in *E. coli* and incubated with SeH4-His or SeH3-His, purified (pull-down) by glutathione sepharose beads. Recombinant GST was control. SeH4-His was pulled down by SeASF1-GST. **E**, SeDJ-1 was cloned into plasmid pGEX-6P-1. Flag-SeASF1 or SeH4 was cloned into plasmid pET28a. SeH4-His and Flag-SeASF1-His were both retained by SeDJ-1-GST. **F**, SeASF1 was cloned into plasmid pDL2, SeH4 or SeH3 was cloned into plasmid pFL7. Total proteins were extracted from *F. graminearum* protoplasts expressing SeASF1-GFP, SeH4-Flag, and SeH3-Flag. Recombinant GFP or Flag was control. The immune complexes were immunoprecipitated with α-Flag antibody (α-Flag IP). Coprecipitation of SeH4-Flag or SeH3-Flag was detected by immunoblotting. **G**, SeH4 was cloned into plasmid pDL2. SeDJ-1 was cloned into plasmid pFL7. Total proteins were extracted from *F. graminearum* protoplasts expressing SeASF1-GFP, SeH4-GFP, and SeDJ-1-Flag. Coprecipitation of SeDJ-1-Flag was detected by immunoblotting. Membranes were stained with Ponceau S to confirm equal loading. Protein sizes are indicated in kDa. Each experiment was repeated at least three times.

### SeDJ-I is involved in PI3K/AKT signaling pathway and interacts with SePI3K or SeGSK3 in *S. eturmiunum*

The PI3K/AKT signaling pathway is a classic signaling cascade that regulates cell growth and proliferation by affecting a multitude of complementary downstream pathways. The previous studies revealed that DJ-1 could increase the AKT phosphorylation and activated the PI3K/AKT signaling pathway in human (Yang et al., 2005; Zhang et al., 2016). However, it has rarely been reported in fungi. Accordingly, the expression levels of *Sepi3k* and *Segsk3* were quantified in two Si*Sedj-1* lines. At the same time, two Si*Sepi3k* lines were obtained using RNA interference. The expression levels of *Sedj-1* and *Segsk3* were examined in two Si*Sepi3k* lines. The results showed that the expression levels of *Sepi3k* or *Sedj-1* were down-regulated in two Si*Sedj-1* or two Si*Sepi3k* lines (Figure 4A), while the expression levels of *Segsk3* was up-regulated in two Si*Sedj-1* lines and two Si*Sepi3k* lines, respectively (Figure 4B). Therefore, *Sedj-1*, a developmental activator, was an upstream component of the *Sepi3k* and *Segsk3* modules that lied in the PI3K/AKT signaling pathway of *S. eturmiunum*. To verify whether SeDJ-1 interacted with these two components in PI3K/AKT signaling pathway, Y2H was first used to ascertain SeDJ-1 interaction with SePI3K or SeGSK3 (Supplemental Figure S20). Subsequently, SeDJ-1 was confirmed to interact with SePI3K or SeGSK3 by pull down (Figure 4C) (The primal western blots of input results are shown in Supplemental Figure S50A, S51A, while the primal western blots of Pull-down results are shown in Supplemental Figure S50B, S51B). Finally, the experiments of both Y2H and pull down were further assured by Co-IP assay (Figure 4D) (The primal western blots of input results are shown in Supplemental Figure S52A, S53A and S54A, while the primal western blots of IP results are shown in Supplemental Figure S52B, S53B). All these data suggest that SeDJ-1 could motivate the major components of the PI3K/AKT signaling pathway to involve in this pathway activity in *S. eturmiunum*.

**Figure 4.**
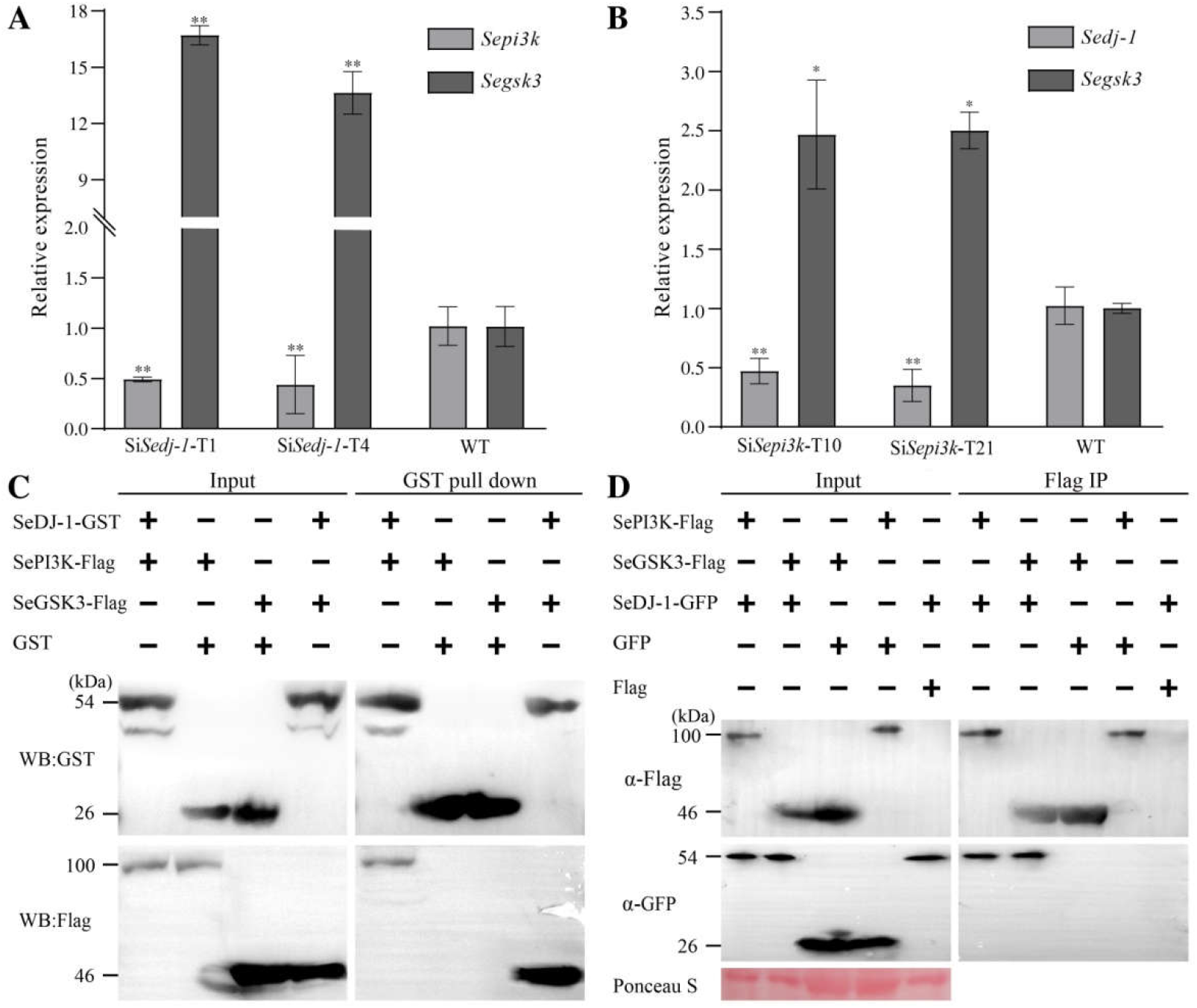
SeDJ-1 is involved in PI3K/AKT signaling pathway and interacts with SePI3K or SeGSK3 in *S. eturmiunum*. **A**, The expression levels of *Sepi3k* and *Segsk3* in two *Sedj-1* lines were measured by qRT-PCR. **B**, The expression levels of *Sedj-1* and *Segsk3* in two Si*Sepi3k* lines were quantified by qRT-PCR. The degree of WT was assigned to value 1.0. Two *Sepi3k*-silenced lines were Si*Sepi3k*-T10 and Si*Sepi3k*-T21. *S. eturmiunum Actin* was used as endogenous control. The bars indicated statistically significant differences (ANOVA; \**P* < 0.05, \*\**P* < 0.01). **C**, SeDJ-1 was cloned into plasmid pGEX-6P-1. Flag-SePI3K or Flag-SeGSK3 was cloned into plasmid pET28a. Recombinant SeDJ-1-GST, Flag-SePI3K-His and Flag-SeGSK3-His were expressed in *E. coli*. SeDJ-1-GST was incubated with Flag-SePI3K-His or Flag-SeGSK3-His and subsequently purified (pull-down) by glutathione sepharose beads. GST-SeDJ-1 was both pulled down Flag-SePI3K-His and Flag-SeGSK3-His. **D**, SeDJ-1 was cloned into plasmid pDL2. SePI3K or SeGSK3 was cloned into plasmid pFL7. Total proteins were extracted from *F. graminearum* protoplasts expressing SeDJ-1-GFP and SePI3K-Flag, SeDJ-1-GFP and SeGSK3-Flag. The immune complexes were immunoprecipitated with α-Flag antibody (α-FLAG IP), and the bound protein was detected by immunoblotting. Membranes were stained with Ponceau S to confirm equal loading. Protein sizes are indicated in kDa. Each experiment was repeated at least three times.

### M6 domain of *Sedj-1* recovers *Sepi3k* silenced transformants to produce perithecia

We previously showed that SeDJ-1 interacted with SePI3K or SeGSK3 and involved in the activity of PI3K/AKT signaling pathway. To decide the critical segment of SeDJ-1 carried out all those functions, seven truncations of *Sedj-1* were obtained and tested whether each of them interacts with SePI3K by Y2H and Co-IP. The results determined that *Sedj-1*-M6 was a key domain for modulating *Sedj-1* interaction with SePI3K (Figure 5A, B) (The primal western blots of input results are shown in Supplemental Figure S55A, S56A and S57A, while the primal western blots of IP results are shown in Supplemental Figure S55B, S56B). Subsequently, *Sedj-1, Sedj-1*-M6, *Sedj-1*-M7 and *Sepi3k* were overexpressed in Si*Sepi3k* strains, respectively. Eight overexpression transformants, OE*Sedj-1* (T13 and T30), OE*Sedj-1*-M6 (T5 and T10), OE*Sedj-1*-M7 (T18 and T26), and OE*Sepi3k* (T8 and T13), were obtained. Those overexpression transformants were identified by western blot and qRT-PCR (Figure 5D, E) (The primal western blots results are shown in Supplemental Figure S58-S61). As a result, these overexpression strains excluded OE*Sedj-1*-M7 (T18 and T26) restored the sexual characteristics in Si*Sepi3k* strains and produced abundant perithecia (Figure 5C). The asexual characteristics of these eight overexpression transformants and two Si*Sepi3k* transformants were unanimous with those of WT strain (Supplemental Figure S22A). The nuclei distributions in mycelia of these transformants were identified with WT strain (Supplemental Figure S22B). All those results indicate that *Sedj-1*-M6 is a key functional domain for modulating sexual features and plays an important role in PI3K/AKT signaling pathway of *S. eturmiunum*.

**Figure 5.**
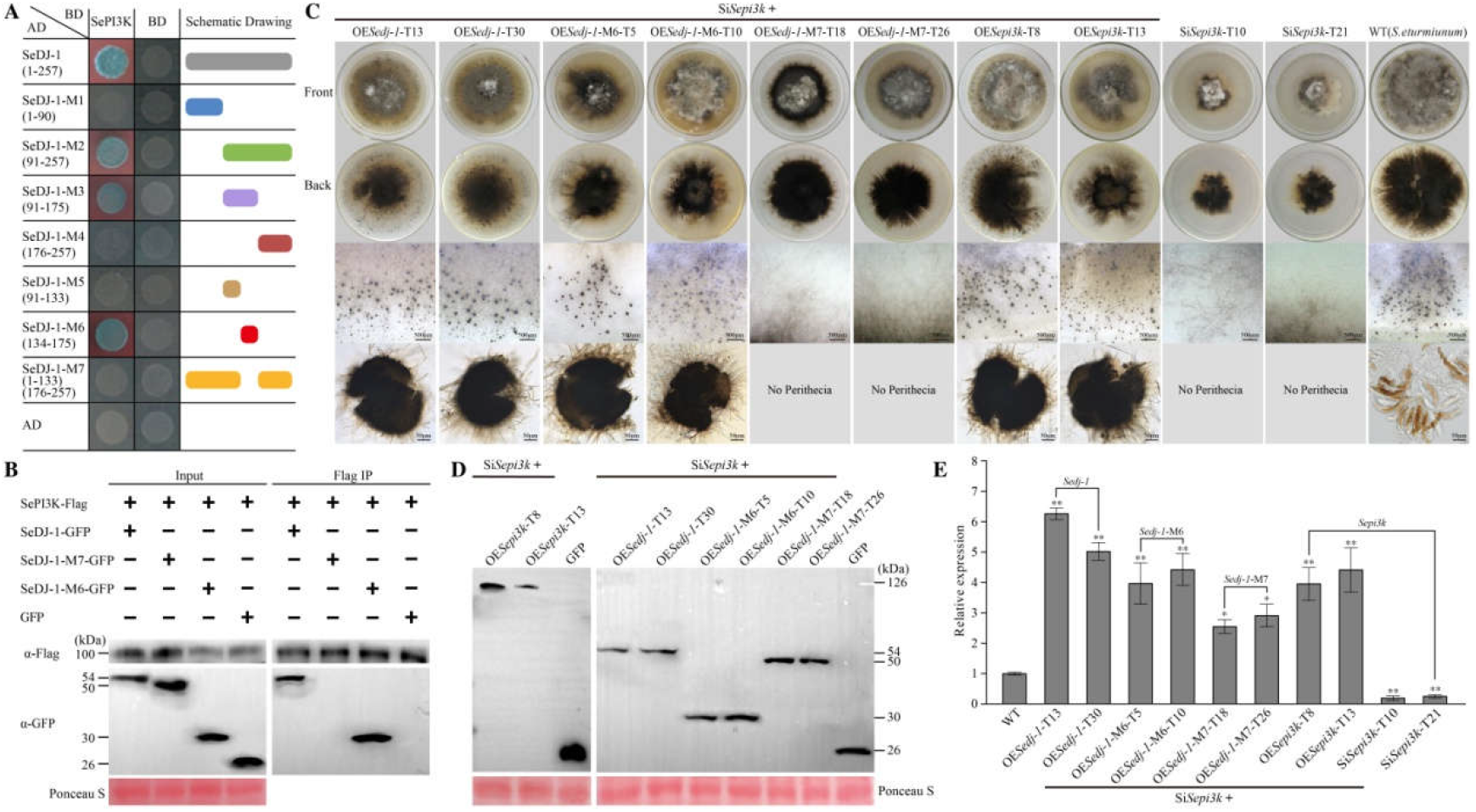
M6 domain of *Sedj-1* induces *Sepi3k* silenced transformants to recover perithecia. **A**, Seven truncations of *Sedj-1* interacted with full length SePI3K by Y2H. **B**, SePI3K was cloned into plasmid pFL7. Three selected truncations, SeDJ-1, SeDJ-1-M6 and SeDJ-1-M7, were cloned into plasmid pDL2, respectively. Total proteins were then extracted from *F. graminearum* protoplasts expressing SePI3K-Flag, SeDJ-1-GFP, SeDJ-1-M6-GFP and SeDJ-1-M7-GFP alone. GFP-fusion was used as control. The immune complexes were immunoprecipitated using α-Flag antibody (Flag IP). Coprecipitation of SePI3K-Flag was detected by immunoblotting. **C**, *Sedj-1, Sedj-1*-M6, *Sedj-1*-M7 and *Sepi3k* were overexpressed in the *Sepi3k* silenced strains, respectively. Eight overexpression transformants, OE*Sedj-1* (T13 and T30), OE*Sedj-1*-M6 (T5 and T10), OESedj-1-M7 (T18 and T26), and OESepi3k (T8 and T13), were obtained and cultured on PDA medium for inducing perithecia production. *Sepi3k* silenced and WT strains were used as controls. The images of perithecia were photographed after growing for 30 days. Those overexpression transformants excluded OE*Sedj-1*-M7 (T18 and T26) produced abundant perithecia compared with Si*Sepi3k* strains. Bar= 50 μm and 500 μm. **D**, Those overexpression transformants were identified by western blot using GFP antibody. **E**, The expression levels of *Sedj-1, Sedj-1*-M6, *Sedj-1*-M7 and *Sepi3k* within corresponding to overexpression transformants were measured by qRT-PCR related to two *Sepi3k* silenced lines. The degree of WT was assigned to value 1.0. *Actin* gene of *S. eturmiunum* was used as endogenous control. The bars indicated statistically significant differences (ANOVA; \**P* < 0.05, \*\**P* < 0.01). Each experiment was repeated at least three times.

**Figure 6.**
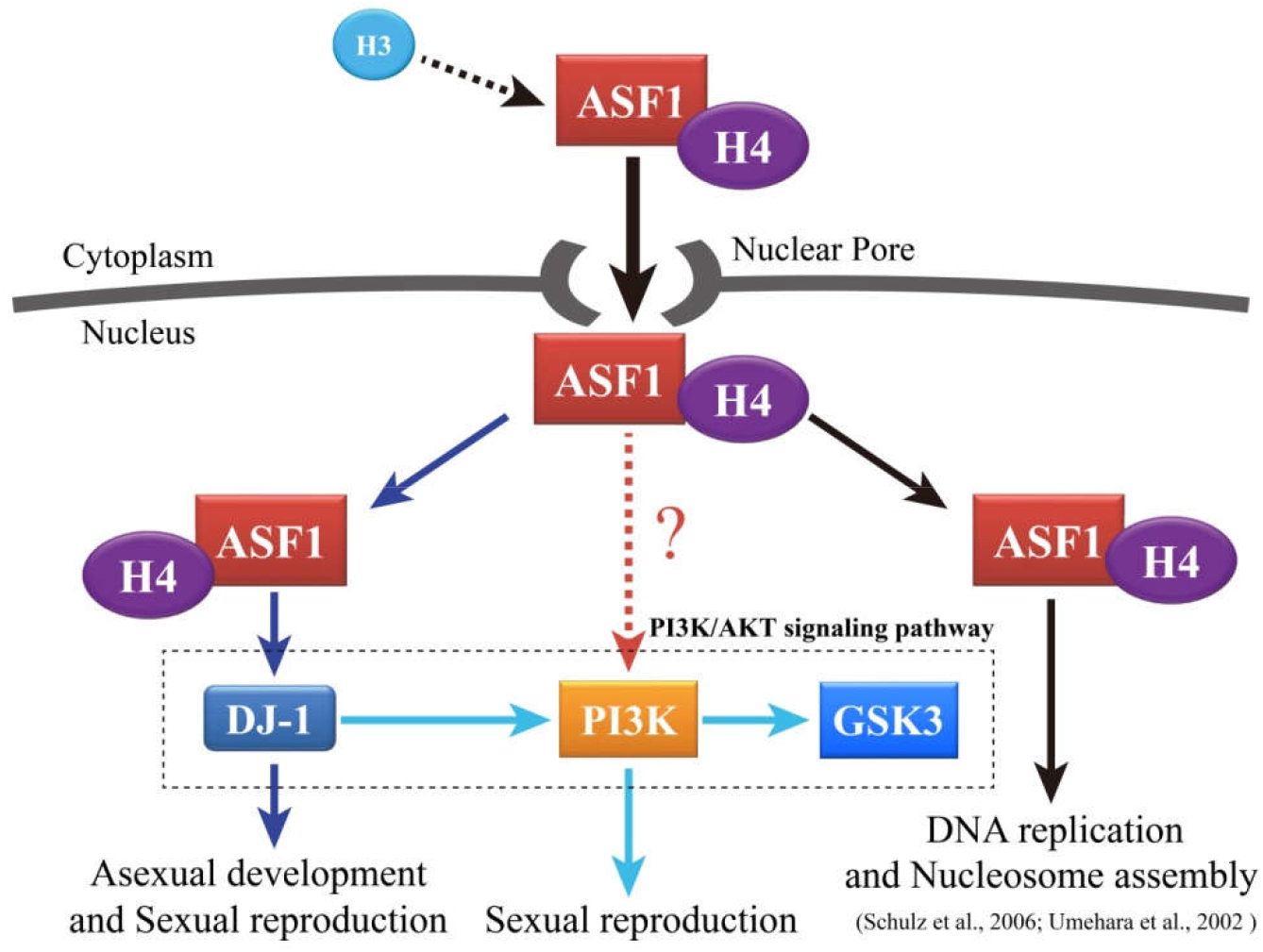
A model for ASF1 binding H4 (ASF1-H4) activates DJ-1 to mediate sexual and asexual reproduction in *S. eturmiunum*. ASF1, a molecular chaperone, interacts with H4 and then translocates into nucleus through the nuclear pore. After getting into nucleus, the dimer of ASF1-H4 modulates DNA replication and nucleosome assembly. ASF1-H4 combines with DJ-1 constituting a new trimeric complex that plays a novel role for modulating sexual and asexual reproduction. Subsequently, DJ-1 also participates in PI3K/AKT signaling pathway for regulating sexual reproduction. Here, it is unknown whether the dimer of ASF1-H4 directly activates PI3K to involve in PI3K/AKT signaling pathway for sexual reproduction process.

### The sexual reproduction of Si*Sedj-1* strains was recovered by overexpressing *Sepi3k*

In our previous study, *Sedj-1* was confirmed to be a positive regulator for effecting on *Sepi3k* mediated sexual development in upstream of PI3K/AKT signaling pathway, we attempted to evaluate whether *Sepi3k* was likely to a reverse regulator for dedicating to *Sedj-1* modulated sexual states. Accordingly, *Sepi3k* and *Sedj-1* were overexpressed in Si*Sedj-1* lines, respectively. Four overexpression transformants, OE*Sedj-1*-T5, OE*Sedj-1*-T20, OE*Sepi3k*-T8, and OE*Sepi3k*-T12, were generated to investigate the role of *Sepi3k* in Si*Sedj-1* strains. Those overexpression transformants were identified by western blot and qRT-PCR (Supplemental Figure S21B, C) (The primal western blots results are shown in Supplemental Figure S62, S63). At 25 days, two OE*Sepi3k* strains similar to two OE*Sedj-1* strains produced abundant perithecia subsequently but Si*Sedj-1* strains did not (Supplemental Figure S21A). However, the asexual characteristics of these four transformants were unanimous with WT strain (Supplemental Figure S22A). The nuclei distributions in mycelia of these transformants were identified with WT strain (Supplemental Figure S22B). Together, these results support our hypothesis that *Sepi3k* regulates sexual reproduction in Si*Sedj-1* strains reversely.

## Discussion

ASF1 was widely present in animals, plants and fungi, but whether ASF1 was related with sexual and asexual reproduction in fungi was scarcely understood. The sexual reproduction of few filamentous fungi species had been described (Coppin et al., 1997). *Stemphylium* is an important genus in filamentous fungi, but typical species *S. eturmiunum* has only one ASF1 (SeASF1) which has more than 90% identify with ASF1 of *S. lycopersici* and other nine fungi species but shares less than 50% similarity with ASF1 of *S. pombe*. SeASF1 carried out the same localizations and sexual reproduction characters as SmASF1 of *S. macrospora* (Gesing et al., 2012). Moreover, SeASF1 could manipulate sexual and asexual reproduction obviously in *S. eturmiunum*, in which transcriptional levels of a serious of genes in MAPK signaling pathway related to sexual reproduction were hardly changed in Se*Δasf1* mutants. Thus, we supposed that SeASF1 was not involved in MAPK signaling pathway related to the fungi sexual mating (Chen et al., 2002; Saito, 2010) and might activate other signaling pathways for sexual mating in filamentous fungi. To further investigate this hypothesis, we performed a transcriptome analysis of SeΔ*asf1*, identifying SeDJ-1 and other six genes (Supplemental Table 4) as possible candidates for cooperating with SeASF1 to modulate the sexual and asexual development. Also, SeDJ-1 plays a crucial role in sexual and asexual development of *S. eturmiunum* in contrast to other six genes by observing phenotypes of each silenced transformant.

Previous studies revealed that DJ-1 was involved in multiple biological functions in mammals (Hijioka et al., 2017; Scumaci et al., 2020; Mencke, 2021; Nakamura et al., 2021). However, DJ-1 regulation of sexual and asexual differentiation in mammals, plants and fungi was poorly understood. Here, we found that SeDJ-1 could involve in sexual and asexual development in *S. eturmiunum*, and further illuminated the mechanisms of SeDJ-1 infiltrating sexual and asexual development. These mechanisms were further supported by transcriptional levels of *Seasf1, SeH4*, and *Sedj-1* in knockout or silenced lines along with SeASF1 interaction with SeH4, and SeDJ-1 interaction with SeASF1 or SeH4 in *vivo* and *vitro*, respectively. Thus, all those results suggested that SeDJ-1 could cooperate with SeASF1 and SeH4 to arouse the sexual and asexual activity.

Sufficient evidences proved that DJ-1 was an important component in PI3K/AKT signaling pathway (Yang et al., 2005) and might bind with various other factors in this pathway to activate a variety of attractive biological processes (van der Brug et al., 2008; Vasseur et al., 2012). By contrast, multiple downstream proteins were reported in PI3K/AKT signaling pathway in mammals (Wang et al., 2013; Xu et al., 2020; Sitaram et al., 2009; Vasseur et al., 2009), but one downstream protein GSK3 (SeGSK3) was found in PI3K/AKT signaling pathway in *S. eturmiunum*. Significantly, SeGSK3 exhibited up-regulated trends in both *Sedj-1* and *Sepi3k* silenced transformants, while SePI3K and SeDJ-1 showed down-regulation in *Sedj-1* or *Sepi3k* silenced transformants, respectively. SeDJ-1 should be upstream component of the SePI3K and SeGSK3 modules in PI3K/AKT signaling pathway of *S. eturmiunum*. Moreover, SeDJ-1 also interacted with SePI3K or SeGSK3 in *vivo* and *vitro*. Therefore, SeDJ-1 should be a key upstream protein in PI3K/AKT signaling pathway and was expected to activate it to regulate sexual and asexual reproduction.

To further investigate whether SeDJ-1 can involve in sexual and asexual reproduction in *S. eturmiunum* and how it does this work. We verified SeDJ-1 and SePI3K could stimulate sexual activity effectively of *S. eturmiunum* in SeDJ-1 or SePI3K silenced strains compared with their overexpression strains, respectively. Meanwhile, SePI3K overexpression in Si*Sedj-1* strains could recover the sexual states of these strains. Thus, SeDJ-1 and SePI3K are not only two important components of the PI3K/AKT signaling pathway, but also carry out the same functions for regulating sexual development in *S. eturmiunum*. SeDJ-1 could interact with SePI3K in our experiments. To further illuminate the mechanism of SeDJ-1 regulating sexual reproduction, the seven truncations of SeDJ-1 were obtained. As a result, SeDJ-1-M6 was defined as a critical segment for interaction of SeDJ-1 with SePI3K and was also proved to be an essential segment for sexual reproduction in OE*Sedj-1*-M6 compared with OE*Sedj-1*-M7. Thus, SeDJ-1-M6 plays a critical role in PI3K/AKT signaling pathway for irritating sexual and asexual reproduction in *S. eturmiunum*. A model is shown in Figure6, SeASF1 coupling SeH4 translocated into the nucleus followed by motivating SeDJ-1 to irritate sexual and asexual means and then SeDJ-1 aroused SePI3K to modulate sexual states. Therefore, SeASF1 could activate PI3K/AKT signaling pathway to regulate sexual and asexual differentiation in filamentous fungi.

## Materials and methods

### Strains and culture conditions

*Stemphylium eturmiunum* strain (EGS 29-099) (WT), *Seasf1* knockout mutants (SeΔ*asf1*), *Seasf1* complemented transformants (SeΔ*asf1*::EGFP*Seasf1*), *Seasf1* heterologous expression transformants (SmΔ*asf1*::EGFP*Seasf1*), *Sedj-1* silenced transformants (Si*Sedj-1*-T1 and Si*Sedj-1*-T4) and overexpression transformants (OE*Sedj-1* and OE*Sepi3k*) were cultured in the dark condition at 25°C on complete medium (CM), or potato dextrose agar (PDA) medium for mycelial growth assays. For the hyphal growth measure, each of these strains was inoculated and grown for 9 days at 25°C in glass dish containing 20 mL PDA medium. To determine the morphology of conidia and ascospores, all of the strains were grown on PDA or CM medium (casein acid hydrolysate 0.5 g/L, casein enzymatic hydrolysate 0.5 g/L, glucose 10.0 g/L, Ca(NO_3_)_2_·4H_2_O 1.0 g/L, KH_2_PO_4_ 0.2 g/L, MgSO_4_·7H_2_O 0.25 g/L, NaCl 0.15 g/L, yeast extract 1.0 g/L, and agar 15.0 g/L) at 25°C. *Escherichia coli* DH5α or *Agrobacterium tumefaciens* AGL-1 was incubated in LB (Luria-Bertani) medium at 37°C or 28°C, respectively (Lennox, 1995).

### Cloning and plasmid construction

Cloning and propagation of recombinant plasmids were done under standard conditions (Sambrook et al., 2001). Deletion of *S. eturmiunum asf1* by homologous recombination was achieved as follows. Briefly, the *Seasf1* flanking regions, 1500 bp upstream and 1500 bp downstream of open reading frame, were amplified using primer pairs *Seasf1*-5f/*Seasf1*-5r and *Seasf1*-3f/*Seasf1*-3r, respectively. The sequences of these primers are summarized in Supplemental Table S2. The upstream fragment was inserted into *Seasf1* knockout vector pXEH by *XhoI*/*BglII*-digested. Then the downstream fragment was inserted into *BamHI*/*XbaI* sites of the vector *Seasf1-*L-pXEH. The vector pXEH carrying a *Hph* resistance cassette was flanked by the *Seasf1* upstream and downstream sequences (Supplemental Figure S5A).

For complementation and heterologous expression analysis, *S. eturmiunum asf1* was cloned from *S. eturmiunum* genome (This genome did not upload) with primers *Seasf1*-pHDT-F/*Seasf1*-pHDT-R, and then cloned into eGPF-pHDT vector. Subsequently, recombinant plasmid eGFP-pHDT-*Seasf1* was transformed into the SeΔ*asf1* mutants and *S. macrospora* Δ*asf1* (SmΔ*asf1*) mutants (S90177) by *A. tumefaciens* mediated transformation (ATMT) method, respectively (Bernardi-Wenzel et al., 2016). Transformants, resistant to G418, were screened by PCR and western blot.

For co-immunoprecipitation (Co-IP) analysis, SeASF1, histone (H3/H4), and SeDJ-1 were amplified from *S. eturmiunum* with primers (Supplemental Table S2), and cloned into the pDL2 or pFL7 in yeast (XK125) by recombination approach (Zhou et al., 2011). Recombinant plasmids were then co-transformed into the protoplasts of *Fusarium graminearum* wild-type strain (PH-1). Transformants were also screened by western blot.

RNA interference (Zhao et al., 2016) was used for *Sedj-1* silencing. The complementary cDNA fragments from *Sedj-1* (499 bp) was amplified from *S. eturmiunum* using primers in Supplemental Table S2 and inserted into vector pCIT that flanked to the intron to form silencing construct, respectively (Zhao et al., 2016). The constructed plasmid pCH-*Sedj-1* was transformed into *S. eturmiunum* strain by *A. tumefaciens* mediated transformation (ATMT) method (Bernardi-Wenzel et al., 2016). For overexpression analysis, *S. eturmiunum Sedj-1* and *Sepi3k* gene were cloned from *S. eturmiunum* strain with primers in Supplemental Table S2, and then cloned into eGFP-pHDT vector, respectively. Subsequently, recombinant plasmid eGFP-pHDT-*Sedj-1* or eGFP-pHDT-*Sepi3k* was transformed into the Si*Sepi3k* lines by ATMT method (Bernardi-Wenzel et al., 2016). Overexpression transformants, resistant to G418 were screened by qRT-PCR and western blot.

### DNA extraction and Southern blot

All strains were inoculated in PDA medium and grown at 25°C for 7 days in the dark condition. Genomic DNA was extracted from mycelia by CTAB (Storchova et al., 2000). Southern blot was performed using the DIG High Prime DNA Labeling and Detection Starter kit I according to the manufacturer’s instructions (Roche Diagnostics, Mannheim, Germany). The specific sequence was amplified from *Hph* gene using primer pairs (Supplemental Table S2), and it was then produced a DIG-labeled probe for hybridization. Each experiment was repeated at least three times.

### RNA extraction and qRT-PCR

Total RNA was extracted from mycelia of *S. eturmiunum* growing in PDB (Potato Dextrose Broth) cultures using the Fungal RNA Kit (OMEGA Biotechnology, USA). Reverse transcription was done using 1 µg of total RNA per 20 µL reaction. SYBR Color qRT-PCR was performed in 20 µL reactions that included 0.4 µg of cDNA, 0.4 µL of gene-specific upstream and downstream primers, 10 µL of 2 × ChamQ SYBR Color qPCR Master Mix (Vazyme) and 5.2 µL of ddH_2_O. The qRT-PCR was performed on an ABI QuantStudio™ 6 Quantitative Real-Time PCR System (Applied Biosystems) under the following conditions: 95°C for 5 min, 40 cycles at 95°C for 10 s, and 60°C for 30 s to calculate cycle threshold values, followed by a dissociation program of 95°C for 15 s, 60°C for 1 min, and 95°C for 15 s to obtain melt curves. Relative expression levels of all above selected genes were determined by qRT-PCR with specific primers listed in the Supplemental Table 2. Changes in the relative expression level of each gene were calculated by the 2^-ΔΔCT^ method (Livak and Schmittgen, 2001). The housekeeping gene *Actin* was used as an internal standard in each case. This experiment was repeated at least three times.

### Gene transcription

For total RNA extraction, WT and SeΔ*asf1* strains were grown for 4 days in PDB medium by the Fungal RNA Kit (OMEGA Biotechnology, USA). The eligible mRNA was enriched with magnetic beads with Oligo (dT). Fragmentation buffer was then added to break the mRNA into short fragments. Using mRNA of WT or SeΔ*asf1* as a template, first-strand cDNA was synthesized with random hexamers, then buffer, dNTPs and DNA polymerase I were added to synthesize second-strand cDNA, followed by using AMPure XP beads to purify double-stranded cDNA. Subsequently, the purified double-stranded cDNA was then subjected to end repair, a tail was added and linked to the sequencing adapter, and then AMPure XP beads were used for fragment size selection. Finally, PCR enrichment was performed to obtain a final cDNA library.

RNA-Seq analysis of total RNA from the WT and SeΔ*asf1* stains was performed by Illumina Hiseq4000 (Berry Genomics, Beijing). Approximately 300 bp fragments were inserted into every library, in which 100 bp sequences were read. Low-quality raw reads were filtered. Resulting paired-end sequencing reads were aligned and quantified using TopHat and Cufflinks with default parameter values. De novo transcriptome analysis was used to estimate transcript abundance and differential expression. Gene expression was calculated as fragments per kilobase of transcript per million mapped fragments (FPKM).

### Gene Ontology

Enriched terms from gene ontology (GO) Biological Process, KEGG (Kyoto Encyclopedia of Genes and Genomes), Swiss-Prot (A manually annotated and reviewed protein sequence database), PIR (Protein Information Resource) and PRF (Protein Research Foundation) databases were identified using the available tools at FungiDB (Stajich et al., 2012) and Blast2GO v2.5. To characterize the genes identified from the differentially expressed genes (DEGs), the GO-based trend tests were performed using the Fisher’s exact test. Fold change > 2.0, *P* value < 0.005 were considered statistically significant.

### Yeast two-hybrid

To test whether SeASF1 and H4 interact with SeDJ-1, Y2H assay was performed according to the Yeast Protocols Handbook (Clontech) using the Y2H Gold yeast reporter strain (Clontech). The *Seasf1, SeH4, SeH3* and *Sedj-1* were amplified by PCR from the cDNA of the *S. eturmiunum*. Then, PCR products were purified and digested with restriction enzyme. The *Seasf1* or *SeH4* was inserted into pGBKT7 plasmid. The *Sedj-1* or *SeH4* was inserted into pGADT7 plasmid. Recombinants of *Seasf1-*BD and *Sedj-1-*AD, *Seasf1-*BD and *SeH4-*AD, *SeH4-*BD and *Sedj-1-*AD were co-transformed into yeast strain Y2H gold, respectively. The transformants were screened on SD/-Trp/-Leu medium (TaKaRa Bio) at 30°C for 3-5 days and assayed for growth on the SD/-Trp/-Leu/-His/-Ade/X-α-gal plates (TaKaRa Bio). Each experiment was repeated at least three times.

### Recombinant protein purification and GST pull-down

The *Seasf1* was cloned into the pET28a vector after adding a 1×FLAG tag to the 5′-terminal of *Seasf1* by PCR. The *Sedj-1* was cloned into the pGEX-6P-1 vector. For the expression of Flag-SeASF1-28a and GST-SeDJ-1, the pET28a construct or pGEX-6P-1 construct was transformed into *E. coli* Transetta (DE3) (Transgene, Beijing, China), and cells were grown to OD_600_=0.6-0.8 at 37°C and then induced with 1M IPTG (isopropyl-β-D-thiogalactoside) for 12-16 h at 16°C. The cells were harvested by centrifugation for 5 min at 8000 rpm at 4°C. The Flag-SeASF1-28a protein cells were resuspended in Ni-lysis buffer (30 mM Tris-HCl, 300 mM NaCl, 30 mM Imidazole, pH 7.5) and lysed with a Ultrasonic Cell Disruptor. The lysate was centrifuged for 30 min at 14 000 rpm (4°C), and the supernatant was passed over a Ni-affinity column (GE) three times at least. Flag-SeASF1-28a was eluted by Ni-elution buffer (30 mM Tris-HCl, 300 mM NaCl, 6 M Imidazole, pH 7.5). The GST-SeDJ-1 protein cells were resuspended in GST-lysis buffer (50 Mm HEPES, 500 mM NaCl, pH 8.0) and lysed with a Ultrasonic Cell Disruptor. The lysate was centrifuged for 30 min at 14 000 rpm (4°C), and the supernatant was passed over a GST-affinity column (glutathione sepharose™ 4B beads GE Healthcare, Little Chalfont, Buckinghamshire, UK) three times at least. GST-SeDJ-1 was eluted by GST-elution buffer (50 mM HEPES, 500 mM NaCl, 10 mM L-glutathione, pH 8.0). The eluent proteins were mixed with loading buffer, and were verified by SDS-PAGE. For glutathione S-transferase (GST) pull-down in vitro, GST-SeDJ-1 and Flag-SeASF1-28a were expressed in *E. coli* strain BL21 (DE3). Total proteins of GST-SeDJ-1 and Flag-SeASF1-28a were then incubated with 4000 μL of glutathione sepharose™ 4B beads at 4°C for 2 h. The supernatant was removed and the beads were washed by GST-lysis buffer three times. Finally, the beads were eluted by GST-elution buffer. Pull-down of GST-SeDJ-1 with Flag-SeASF1-28a was detected using an anti-Flag (Invitrogen). Each experiment was repeated at least three times.

### Co-IP

*F. graminearum* protoplasts were transfected with the indicated combination plasmids and empty construct. Total mycelium proteins of *F. graminearum* were extracted with an extraction buffer [50 mM HEPES, 130 mM NaCl, 10% glycerin, protease inhibitors (25 mM Glycerol phosphate, 1 mM Sodium orthovanada, 100 mM PMSF), pH 7.4]. For FLAG IP, protein extracts were incubated with 30 μL of Anti-Flag^®^ M2 Affinity Gel beads (Sigma-Aldrich) at 4°C for 4 h, The beads were then collected by centrifugation at 3000×*g* and washed five times with a washing buffer (50 mM HEPES, 130 mM NaCl, 10% glycerin, pH 7.4). The bound proteins were eluted from the beads by boiling for 15 min. The beads were collected by centrifugation at 3000×*g* for 2 min. Proteins were separated by 12% SDS–PAGE gels and detected using immunoblotting with a monoclonal α-Flag antibody (Sigma-Aldrich) or a α-GFP antibody (Invitrogen). Membranes were stained with Ponceau solution (CWBIO, China). Each experiment was repeated at least three times.

### Western blot

Protein samples were separated by 12% SDS-PAGE gels at 100 V for 3 h in running buffer (25 mM Tris-base, 200 mM Glycine, 0.1% SDS). Gels were transferred to Immobilon^®^-P PVDF membrane for 1.5 h at 230 mA. Membranes were then blocked in 5% non-fat milk in 1×TBST (0.02M Tris-base, 0.14M NaCl, pH 7.4) with 0.1% (vol/vol) Tween-20 prior to addition of GFP or FLAG antibodies (Sigma-Aldrich) at 1:5000 dilution and incubated at room temperature for 1-1.5h. The membranes were washed three times with TBST and then were incubated for 1h with a horseradish peroxidase labeled immunoglobulin G (IgG-HRP) secondary antibody (Thermo Fisher Scientific, no. 31430) at 1:7500 dilution. The specific proteins were visualized by using the ECL Chemiluminescence Detection Kit (Vazyme). The images were caught by Tanon-5200 Chemiluminescent Imaging System (Tanon, China). Each experiment was repeated at least three times.

### Microscopy

To observe the morphology of conidia and conidiophores, all the transformants and WT strains were grown in the dark condition at 25°C for 4 weeks on PDA medium by inserting double slides. Microscopic examination of nuclear distributions in mycelia, the transformants and WT strains were stained using 4,6-diamidino-2-phenylindole (DAPI). To image the sexual structures including perithecia and asci, all these test strains were cultured on CM medium at 25°C for 6 weeks in dark condition. Perithecia were sectioned by using a double-edged blade in a dissecting microscope (Olympus, SZX10). The asci, conidia and conidiophores were all captured with 20 × or 40 × objectives of Olympus microscope (Olympus BX53, Tokyo, Japan) using differential interference contrast (DIC) and fluorescence illumination. Microscopic characters of asexual structures were further determined by measurements of 50 mature conidia and 50 conidiophores. The experiment was repeated at least three times.

### Statistical analysis

Data were analyzed using Systat 12 (Systat Software Inc., San Jose, CA, USA). The data were subjected to one-way analysis of variance (ANOVA). Student’s t-test was used for two means, and Duncan’s multiple range test of least significant difference (LSD) was used for more than two means. *P* values of 0.05 and 0.01 were used as indicated.

## Supplemental data

**Supplemental Figure S1**. Phylogenetic analysis of ASF1 sequences from *S. eturmiunum* and all other fungi, plants and animals species.

**Supplemental Figure S2**. Alignment of SeASF1 sequence with its homologous sequences from other fungi, plants and animals species.

**Supplemental Figure S3**. Phenotypic characterization of the *Seasf1* heterologous expression in SmΔ*asf1*.

**Supplemental Figure S4**. Heterologous expression transformants of *Seasf1* were verified by PCR and western blot.

**Supplemental Figure S5**. Deletion of the *Seasf1* gene in *S. eturmiunum*.

**Supplemental Figure S6**. Two complemented transformants of *Seasf1* were verified by PCR and western blot contrast to deleted mutants.

**Supplemental Figure S7**. Hyphal and colonial growth of *S. eturmiunum asf1* deleted mutants and complemented transformants.

**Supplemental Figure S8**. The effect of *Seasf1* on the transcriptional regulation of regulators on sexual reproduction.

**Supplemental Figure S9**. Transcriptome analysis of the differentially expressed genes (DEGs) in a SeΔ*asf1* mutant compared with WT-vegetative and WT-sexual strains.

**Supplemental Figure S10**. Transcriptome analysis of the differentially expressed genes (DEGs) in SeΔ*asf1* mutant and two WT strains.

**Supplemental Figure S11**. Phylogenetic analysis of DJ-1 sequences from *S. eturmiunum* and all other fungi, plants and animals.

**Supplemental Figure S12**. The colonial phenotypes of *Sedj-1* silenced transformants.

**Supplemental Figure S13**. The asexual and sexual development in Se01950 silenced transformants.

**Supplemental Figure S14**. The asexual and sexual development in Se03485 silenced transformants.

**Supplemental Figure S15**. The asexual and sexual development in Se04320 silenced transformants.

**Supplemental Figure S16**. The asexual and sexual development in Se07693 silenced transformants.

**Supplemental Figure S17**. The asexual and sexual development in Se10206 silenced transformants.

**Supplemental Figure S18**. The asexual and sexual development in Se10302 silenced transformants.

**Supplemental Figure S19**. SeASF1 interaction with SeH4 or SeDJ-1 and SeH4 interaction with SeDJ-1.

**Supplemental Figure S20**. SeDJ-1 interaction with SePI3K or SeGSK3.

**Supplemental Figure S21**. The sexual reproduction of Si*Sedj-1* strains was recovered by overexpressing *Sepi3k*

**Supplemental Figure S22**. The asexual development of Si*Sepi3k*, and OE*Sedj-1*, OE*Sedj-1*-M6, OE*Sedj-1*-M7 or OE*Sepi3k* in Si*Sepi3k*, and OE*Sedj-1* or OE*Sepi3k* in Si*Sedj-1* transformants.

**Supplemental Table S1**. The ASF1 genes of *Stemphylium eturmiunum* and other organism species.

**Supplemental Table S2**. Primers used in this study.

**Supplemental Table S3**. Plasmids used in this study.

**Supplemental Table S4**. The differentially expressed genes in SeΔ*asf1* vs WT-sexual.

**Supplemental Table S5**. The DJ-1 genes of *Stemphylium eturmiunum* and other organism species.

## Acknowledgements

We thank Minou Nowrousian (Ruhr-Universität Bochum) for providing *S. macrospora* strains. We thank Jingze Zhang (Zhejiang University) for transcriptome analysis and Daohong Jiang (Huazhong Agricultural University) for providing the plasmid. This work was supported by grants from the National Natural Science Foundation of China (31230001, U200220015).

## Author contributions

S.W., Z.L. and X.G.Z. designed the experiments and wrote the paper. S.W., X.L., C.X., S.G., W.X., L.Z. and C.S. performed the experiments. S.W., X.L., C.X., S.G., W.X., Z.L. and X.G.Z. contributed to the data analysis.

